# Melt into the group Electrophysiological Evidence of Gestalt Perception of Human Dyad

**DOI:** 10.1101/2020.11.14.382481

**Authors:** Karima Mersad, Céline Caristan

## Abstract

It has been shown recently that the human brain has dedicated networks for perception of human bodies in synchronous motion or in situation of interaction. However, below motion and interaction, how does the brain process a simple plurality of humans in close positioning? We used EEG frequency tagging technique to investigate integration of human dyad elements in a global percept. We presented to participants images of two silhouettes, a man and a woman flickering at different frequencies (5.88 vs.7.14Hz). Clear response at these stimulation frequencies reflected response to dyad parts, both when the dyad was presented upright and inverted. However, an emerging intermodulation component (7.14 + 5.88 = 13.02 Hz), a nonlinear response regarded as an objective signature of holistic representation, was significantly enhanced in upright relatively to inverted position. Inversion effect was significant only for the intermodulation component as opposed to stimulation frequencies revealing that dyad configuration perception overrides structural properties of dyad elements. Inversion effect was not significant for a pair of non-human objects. Our results show that merely facing two humans in close positioning leads to perceptually bind them and suggest that the perception of individuals is of different nature when they form a plurality.

## Introduction

As intuitive social animals, all of us have noticed that an individual alone and the same individual in a group are not the same psychological beings. Yet, research in social psychology has revealed how important the influence of others is on our behavior. In Solomon Asch’s seminal study on social conformity, participants often went with the majority (actually partners of the experimenter responding according to a prearranged plan) giving the same wrong answer in a trivial visual-perception task (Asch, 1955). Other people in our surroundings do not need to be active in order to have an influence on our minds and behaviors Indeed, in his work on social facilitation (Zajonc, 1965), the author showed that the mere presence of conspecifics peers would increase the performance in certain tasks and decrease the level of accomplishment on other tasks. This effect on the group upon individuals likely reflects sensitivity to complex and subtle dynamics of human aggregate. It is likely to have its roots in specific perceptual mechanisms, where vision could be the most important apparatus (Nakayama et al., 2010). Behavioral and brain-imaging studies have started to characterize, in particular, brain mechanisms dedicated to the perception of individuals engaged in social interaction (Abassi & Papeo, 2020; Isik et al., 2017; Papeo et al., 2017; Walbrin et al., 2018). It is remarkable that this set of experimental research presupposes that a plurality of individuals is essentially represented as the result of explicit interactions between those individuals. An alternative to this view is that the human brain tends to process any plurality of individuals who are merely close together in space as a coherent unit based on a global configuration. Little is known about the process of human group perception, independently of interaction phenomenon, when an individual merely faces a plurality of conspecifics.

As physical entities, humans are visually perceived as faces and/or bodies. Faces and bodies are very specific stimuli of the environment. They are present from the very first moments of life and all along the development of human beings. These singular animate objects are critical for our life and even for our survival: they provide information, knowledge, trigger emotions and might be related to what the philosopher Spinoza calls *conatus*, that is “the effort by which each thing, as far as it can by its own power, strives to persevere in its being”. Unsurprisingly, evolution has endowed human with highly specialized perceptual visual systems in order to process human faces and bodies differently than other objects. For example, when present in a visual scene, faces are immediately detected as they automatically elicit rapid saccades towards them (Crouzet et al., 2010). The human brain will respond with a specific electrophysiological signature when processing this specific visual object (Bentin et al., 1996; Bötzel et al., 1995; Eimer, 2011) and selectively activate dedicated brain networks localized in the right hemisphere (Rossion et al., 2012). Similarly, the posture of human bodies, silhouettes or perception of movement trigger specific behavioral - and cerebral mechanisms. For example, adults identify the biological motion of conspecifics like walking in few hundred milliseconds by the mere observation of a few point-light displayed on the joints of the walker (Johansson, 1973, 1976). This perceptual expertise is detected by neuroimaging studies in adults revealing the existence of brain regions dedicated to body perception. For example, in a study (Jiang et al., 2001), the author showed that adults selectively activate the lateral occipital area of the right hemisphere when perceiving human bodies contrary to the perception of other types of objects.

What are the characteristics of the mechanism underpinning face and body perception? In a seminal publication, Young et al. (1987) brought insight to this question. They showed that when aligning the upper half part of a face (A) with the lower half part of another (B), the resulting composite face results in the percept of a new face. Most interestingly, this new face is perceived coherent enough to impair the upper half of face A recognition as belonging to the face A. These results have been interpreted by the authors as a demonstration that the human brain do not process face component features locally and as separate parts but rather integrate them in a strongly coherent percept based on their global configuration (distance and relative position between parts). This interpretation is supported by the finding that the perception of the composite face as a whole is impaired when the faces are presented upside-down (Young et al., 1987). Indeed, the inversion effect that is decreased accuracy and longer reaction times in face recognition when faces are upside down is typically considered as an evidence for an holistic or a configural processing. What is of major importance for our purpose, Reed et al., (2006) used the inversion paradigm in order to provide the evidence that the human body perception phenomenon also involves a configural processing and the computation of relations among body components (arms, feets etc..).

Nevertheless, recent work has revealed the presence of a distinctive mechanisms and a cortical specialization in the perception of a group of two humans (a dyad) in interaction. In their study Papeo et al., (2017) showed that inversion impairs the detection of facing dyads (two persons – faces and bodies – positioned face-to-face seemingly interacting) more than non-facing dyad. Moreover, facing dyads have been found to draw more attention than non-facing dyads as showed by Papeo & Abassi, (2019) using a visual search task. Finally, compared with non-facing bodies, facing bodies increase activity in specific areas of the visual cortex (Abassi & Papeo, 2020). These behavioral and brain-imaging methods provide evidence for a configural processing of a group of (two, facing) humans. However, they do not provide any answer to an important question: does the presence of a group configuration modifies the perception of individuals that form the group, merely by the fact of their presence inside a group?

Multi-input frequency tagging electrophysiological technique has been recently used to probe configural processing while dissociating the activity underlying the response to whole configuration from the response to local elements forming the image. In this paradigm, different parts of the visual stimuli are presented at different frequencies (for example, two images presented at f1 and f2 frequencies). In addition, to generate steady state visual evoked potentials (SSVEP) at f1, f2 and at their harmonics (nf1 and mf2, m and n being integer numbers), this method allows to capture additional components, known as intermodulation terms (IM). These components are not present in the input and result from non-linear interaction between populations of neurons responding to the image parts (two, in this example). IM responses occur at frequencies that are sums and differences of the different harmonics (nf1 ± mf2).

Frequency Tagging technique is becoming an important tool for studying human cognition, providing objective signatures of neural processes underlying cognitive functions. Using this method Boremanse et al., (2014) brought experimental evidence of an objective neural signature of perceptual binding of face parts. Benefitting from the same technique Alp et al., (2016) could predict precisely and subsequently demonstrate the presence of neural processes specifically involved in the emergence of illusory surface perception (a Kanisza square). More recently Radtke et al., (2020) used Frequency Tagging method to show that the extraction of the meaning of a scene (Gist perception) is correlated with neural association between simultaneously presented objects sharing semantical relation.

In the present study, we aimed at targeting the neural process that takes place during the perception of a group of humans in its simplest form: two individuals positioned close to each other. Our hypothesis is that dyad, just like individual faces and bodies, is processed as a visually structure unit, even when there is no apparent interaction between its parts. To assess the emergence of Gestalt-like properties resulting from the perception of a dyad, we recorded EEG response while simultaneously presenting two human silhouettes tagged at different frequencies. We intended to dissociate objectively the response to elements forming the dyad from the response to the dyad as a whole entity. To better investigate the social character of the holistic representation, we also presented pairs of chairs as non-human control stimuli. We expected the perception of individual human silhouettes (i.e. parts of the dyad) and individual chairs (parts of the pair) to give rise to EEG response at the precise two fundamental frequencies and their harmonics (e.g., f1, f2, 2f1, 2f2, 3f1,..). While perceiving two human beings next to each other tends to evoke a group configuration with a social meaning, perceiving two chairs one beside the other should not evoke the same social meaning. Consequently, we expected the perception of dyad stimulus to trigger a higher response in IM frequencies (f1+f2, f1-f2, 2f1+f2, etc) compared to the perception of the chairs. As the mechanism expected goes beyond the simple pairings of objects belonging to the same semantic category, involving a holistic perception, we also presented inverted silhouettes of dyad and of pair of chairs. We predicted that the response to the global dyad configuration at IM components would be reduced by inversion but not the response to individual silhouettes at stimulation frequencies (Jacques & Rossion, 2009; Suzuki & Cavanagh, 1995).

## Method

### Participants

*30 adults (three left-handed; 17 females; age range: 18–43, mean age 20*.*09 with normal or corrected-to-normal vision participated in the study. 4 additional participants were tested but their data were not included in the analysis for the following reasons: The results of 2 participants were excluded due to the presence of continuous noise in their recording making the data unexploitable whereas the results of 2 participants were excluded due to an error made by the experimenter during the recording. A written informed consent approved by the Research Ethical Committee of the University of Paris was obtained from all participants prior to their participation in the experiment*.

### Stimuli

Two black silhouettes, a man and a woman (head and shoulders) and two black chairs were used as central displays on a white background. Silhouettes, known to induce strong inversion effect (Stein et al., 2012) were used instead of pictures to avoid processing of local face features. The image of the 2 chairs appeared different one from the other in order to be associated with a female and a male category respectively. For example, the ‘female’ chair was slightly thinner (see Figure 1). Stimuli were displayed as pairs of humans and chairs presented upward or upside-down. To facilitate perceptual binding of silhouettes constituting the pair, silhouettes identity was held constant and their position, slightly leaning to each other intended to evoke closeness, in upward as well as in inverted position. Silhouettes and chairs pictures subtended approximately the following dimensions of visual angles: Man silhouette: 7.77×7.67°, Woman silhouette: 7.20×6.79°, Female chair: 5.85×7.69° Male chair: 6.21×7.69°. Distances between stimuli were controlled between human dyad and chair pairs: mean distance between the two closest points: 1.61° of visual angle for both type of stimuli. At the center of silhouette pairs, a black cross (0.3×0.1°) was displayed at random times at the relative same location (middle of the faces and upper chair parts) in upright and inverted condition. The selection of chairs as control stimuli was inspired by a recent study showing a stronger inversion effect for pairs of bodies than for pairs of chairs (Papeo et al., 2017). Care has been taken for stimuli selection. Firstly, to maximize signal-to-noise ratio, the fundamental frequencies and the sum intermodulation (IM) component (5.88 Hz + 7.14 Hz = 13.02 Hz) were located outside of the alpha (8–12 Hz) range (Regan, 1989). In addition, both 5.88 and 7.14 were integer divisions of the 1000 Hz refresh rate. Finally, stimulation frequencies were selected in the (4-9 Hz) theta-band interval reported to elicit maximal responses to contrast-modulated stimulation with social stimuli like faces (Boremanse et al., 2014).

**Figure 1.**
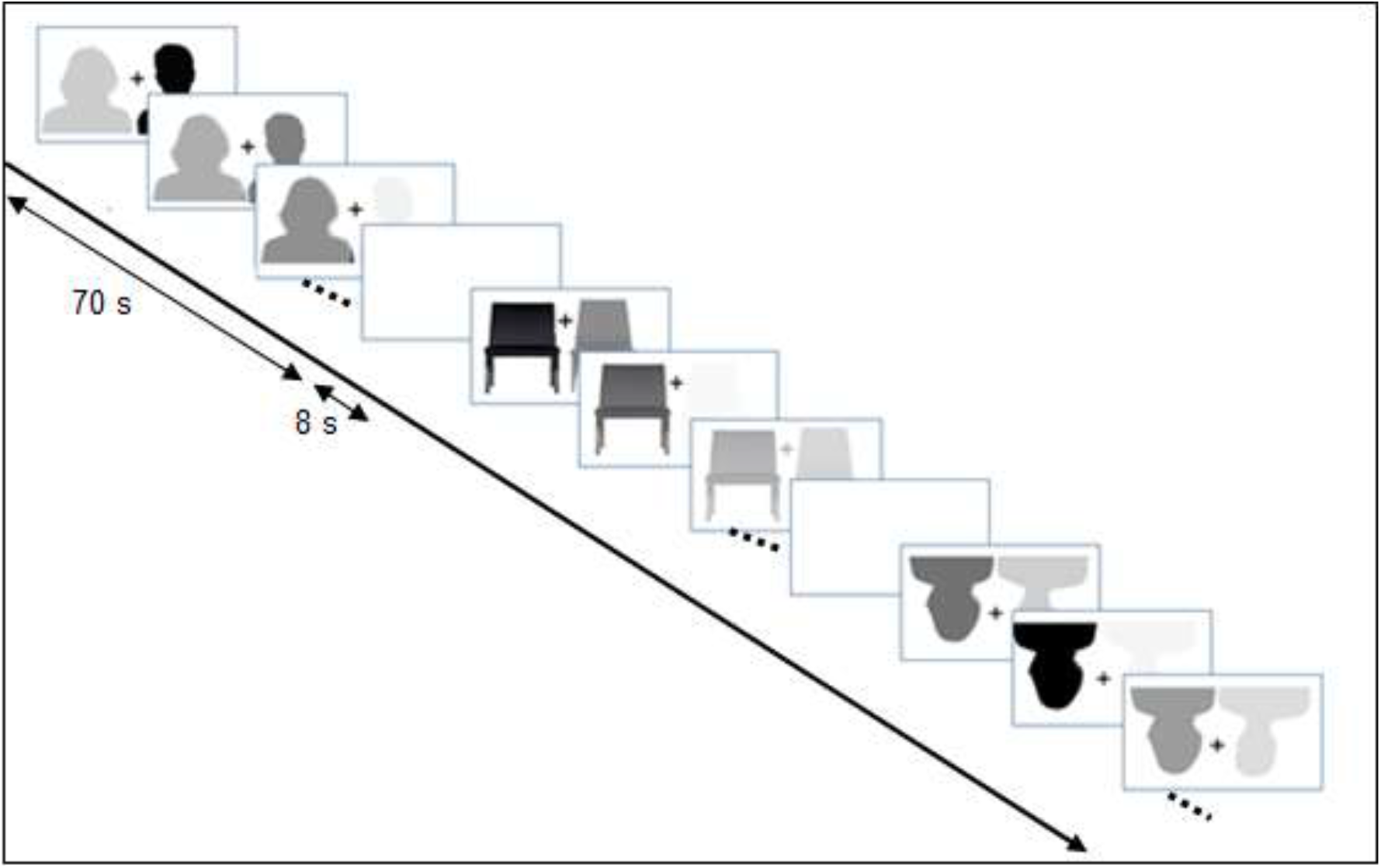
Example of an experiment timecourse. In this example a trial is represented with 3 screen captures. In each trial one pair of stimulus randomly chosen from {Chairs Up, Chairs Inverted, Humans Up, Humans inverted}, was presented during 70 s while each element of the pair was contrast modulated randomly either at f1 or f2. Two trials were separated by a white screen during 8 s.

### Procedure

After electrode-cap placement, participants were seated in a dimly lit room at a distance of approximately 70 cm of the sceen. Sitting chair was adjusted so that the eyes of the participant were in the same level as with the center of the screen. Stimuli were displayed using an in-house application written in Python Psychopy software (Peirce et al., 2019). Figure 1 represents an example of trial time course. Each trial lasted 70 s during which participants viewed pairs of stimuli contrast modulated respectively at f1 and f2. Each participant viewed a total of 12 trials including 4 trials where human silhouettes were presented in upward position, 4 trials where human silhouettes were presented in Inverted position, 2 trials where chair silhouettes where presented in upward position and 2 trials where chair silhouettes were presented in Inverted position. Each human silhouette was alternatively flickering at f1 or f2 and was alternatively appearing on the left or right side in a random and counterbalanced fashion. Each chair was alternatively appearing on the left or right side in a randomized and counterbalanced fashion. Note that unlike (target) Human stimuli, (control) Chair stimuli were not counterbalanced for flickering frequency (a fixed frequency was associated to each chair). This resulted in less trials in Chair than in Human condition. Our rationale was to reduce the fatigue and discomfort driven by the cumulative flickering throughout the experiment and maintain the quality of the recorded data. Two trials were separated by a white screen during 8 s and a break was proposed after 6 trials. The order of presentation of trials was randomized across participants. To help maintaining attention throughout the duration of the trial, participants had to press the space bar whenever the cross was appearing or disappearing at random times at the center of the screen, between the two silhouettes. EEG activity was recorded using a Brainvison amplifier system (Brain Products) with 64 electrodes referenced to the vertex. EEG was digitized at a 1000 Hz sampling rate. To record a highly precise timing of trial onset, triggers were sent from the parallel port of the stimulation computer to the amplifier.

### Data analysis

EEG analysis was conducted using custom-made MATLAB scripts and EEGLAB toolbox (Delorme & Makeig, 2004). The data were resampled to 250 Hz to increase the speed of data processing by reducing the workload. Data were band-pass filtered between 0.1 and 100 Hz so as to remove slow drifts and very high frequencies. After visual rejection of paroxysmal portions of the continuous recording, data were segmented into windows of 66.67 s comprising exactly 392 cycles of f1 (5.88 Hz) and 476 cycles of f2 (7.14). Our purpose in choosing a long duration epoch (66.67 s) was to obtain an amplitude spectrum with a high-frequency resolution (0.015 Hz = 1/66.67). We then removed the first 2 seconds of the recording to avoid artifacts caused by the onset of visual stimulation and to give time to the brain to be entrained by the stimulation. In a following step, we performed automatic rejection of epochs and channels contaminated by artifacts. More precisely, epochs were considered unsuitable for analysis if their fast average amplitude exceeded 250 μV or their deviation between fast and slow running averages exceeding 150 μV. For each subject, channels that had more than 50% of epochs marked as unsuitable were considered as bad channels. 3.4 % of channels on average were rejected per subject. Trials having more than 80% of bad channels were rejected from the analysis. On average, 6,3 % of epochs by subject were rejected. For each participant, epochs of the same condition were averaged to increase the signal to noise ration of the EEG response. We obtained 4 conditions per participant: HumanUp (Human silhouettes in upside position); HumanInvert (Human silhouettes in inverted position), ChairUp (Chair silhouettes in upside position), ChairInvert (Chair silhouettes in inverted position). The resulting averaged waveforms were then submitted to a Fourier decomposition. For each frequency component, the amplitude was taken as the magnitude of the complex number resulting from the FFT for this frequency. Figure 2 shows the amplitude spectrum describing the contribution of each frequency and their topographical maps. SNR of each frequency was then computed as its z-score. More precisely, for each participant, each condition and each electrode, SNR was computed as the amplitude of the signal at the frequency of interest divided by the mean of the amplitude of the 30 neighboring frequency bins (15 from both sides), excluding two bins (one from both sides) adjacent to the bin of interest (Rossion & Boremanse, 2011; Srinivasan et al., 1999). To determine which frequency was above noise level, we computed average of frequency SNR of all participants, all electrodes and all conditions. Five frequencies prove to be above noise threshold (z > 2, p < 0.01): stimulation frequencies: f1 and f2, two harmonics 2f1 and 3f2 and the second order sum IM component: f1+f2. To perform statistical analysis we selected adjacent electrodes with a maximal SNR for the fundamental frequencies, as classically carried out in similar paradigms (e.g., Alp et al., 2016, 2017). 12 electrodes located over parieto-occipital region were selected: CP1, CPZ, CP2, CP4, P1, PZ, P2, P4, POz, PO4, PO8, Oz, O2 (white circles on the maps in Figure 3). Figure 3 depicts SNR frequency spectrum, computed on the region of interest, separately for average Human and Chair conditions in both up and inverted position presentation. A repeated measure analysis of variance (ANOVA) was computed on the same region, with SNR as dependent variable and two factors: Position (Up, Inverted) and Object (Humans, Chairs). We analyzed separately both sets of frequencies selected above noise threshold: on the one hand, stimulation frequencies and their harmonics, on the other hand, IM component.

**Figure 2.**
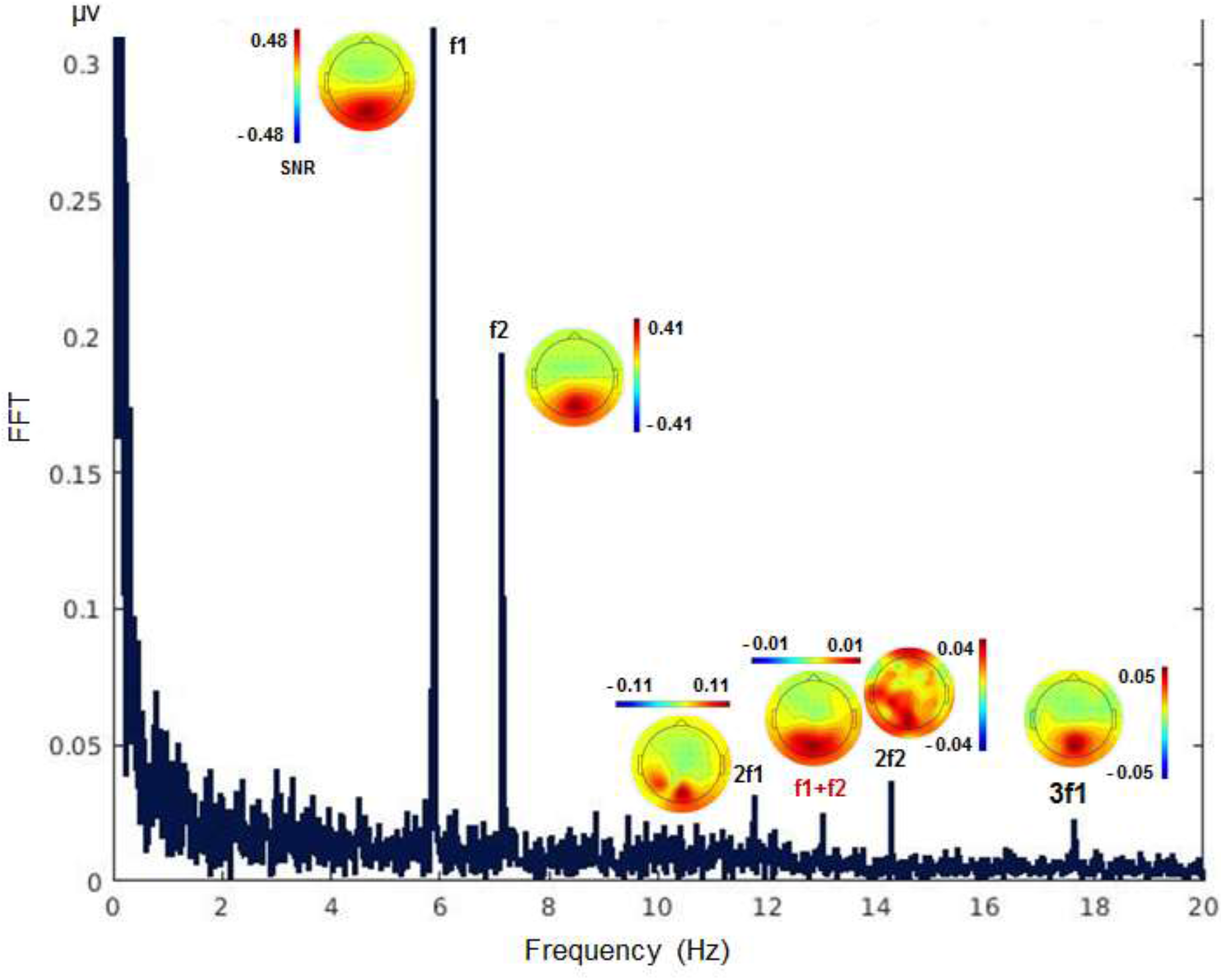
Amplitude Spectrum describing each component frequency contribution (0 – 20 Hz) on average on all participants and all conditions, as well as topographical maps of stimulation frequencies, three harmonics and the second order sum IM component f1+f2 =13.02.

**Figure 3.**
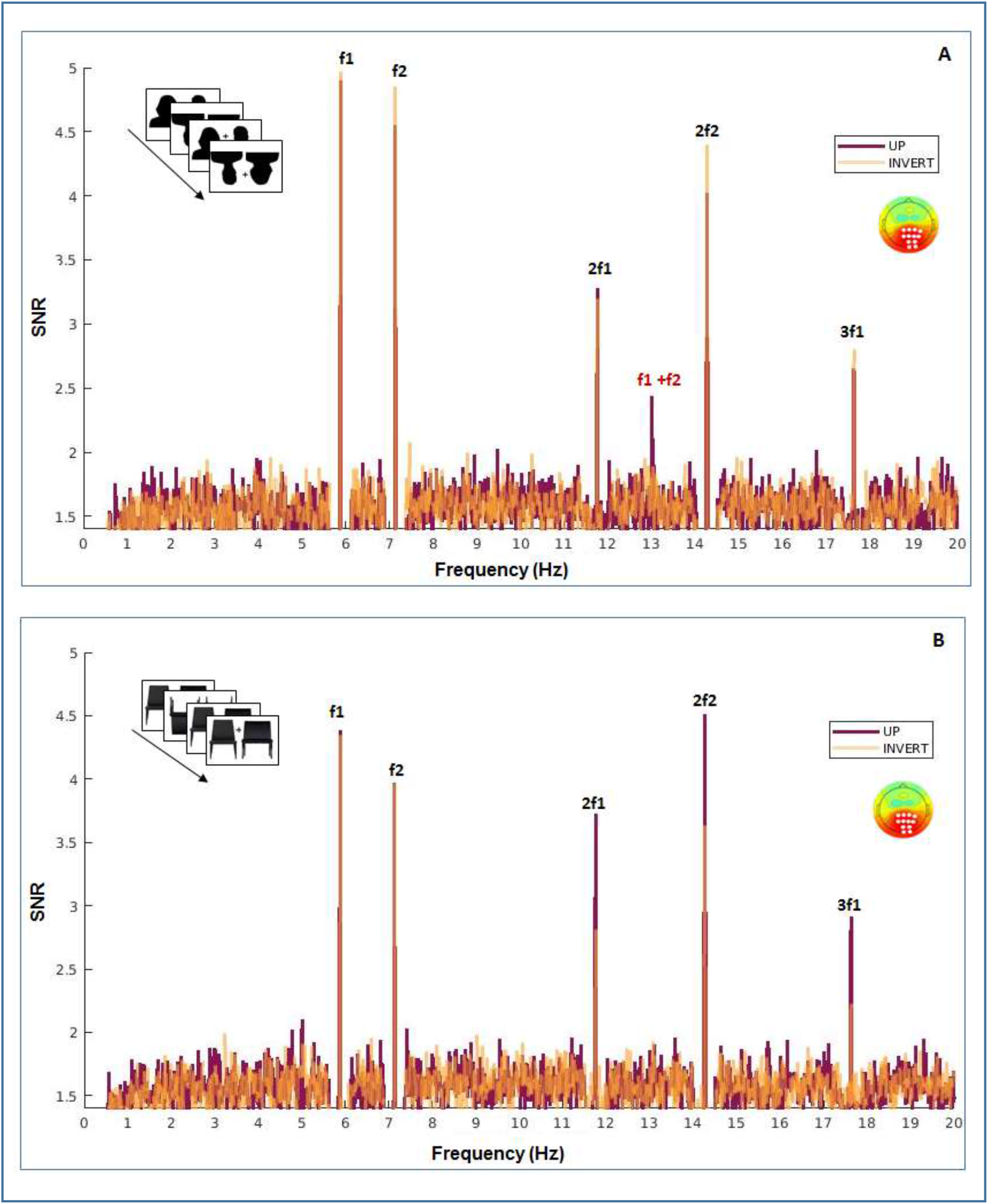
Signal to Noise Ration (0-20 Hz) averaged across participants and cortical region of analysis (white dots on topographical map) for Humans (A) and Chairs (B) conditions. Up and Inverted trials are represented on both figures, with different colors, in transparency. Clear response appears at stimulation frequencies and harmonics in both conditions. The peak at the second order sum IM component (f1+f2 =13.02), present in Humans condition, is higher for Up than Inverted trials and is absent from Chairs condition.

## Results

As can be observed on Figure 2, Amplitude spectrum averaged on all participants, all conditions and all electrodes shows a clear response at the two stimulation frequencies (f1= 5.88 Hz, and f2 = 7.14 Hz) and their harmonics (2f1= 11.76 Hz, 2f2 = 14.28, 3f1= 17.64 Hz, 3f2 = 21.42). As depicted on topographical maps, the response at stimulation frequencies and harmonics was mainly located in occipital region. We observe a typical decrease of spectral amplitude with frequency, nevertheless maintaining second and third order harmonics clearly above noise. Most importantly for our hypothesis, another spectral amplitude peak in the same region is observed at the frequency f1+ f2 (5.88 + 7.14 = 13.02) corresponding to the second order intermodulation frequency. We used SNR values to statistically evaluate and compare stimulation and IM frequencies in the different conditions. To do so, we averaged SNR of both stimulation frequencies f1 and f2, in the one hand, and stimulation frequencies and their second harmonics (the results were similar when including also the third harmonic in the average), on the other hand.

### Response at stimulation frequencies and their harmonics

#### First harmonics (stimulation frequencies)

For the mean f1 and f2 SNR, we observed a significant main effect of Object (F (1, 29) = 25.64, p < 0.001, η^2^ = 0.46), with a higher SNR for Human stimuli (M = 4.82, SD = 0.89) than for Chair stimuli (M = 4.17, SD =0.76), effect size d = 0.79. We observed no main effect of position (F (1, 29) < 1) nor interaction between the two factors F (1.29) < 1. Figure 4 depicts SNR by condition.

**Figure 4.**
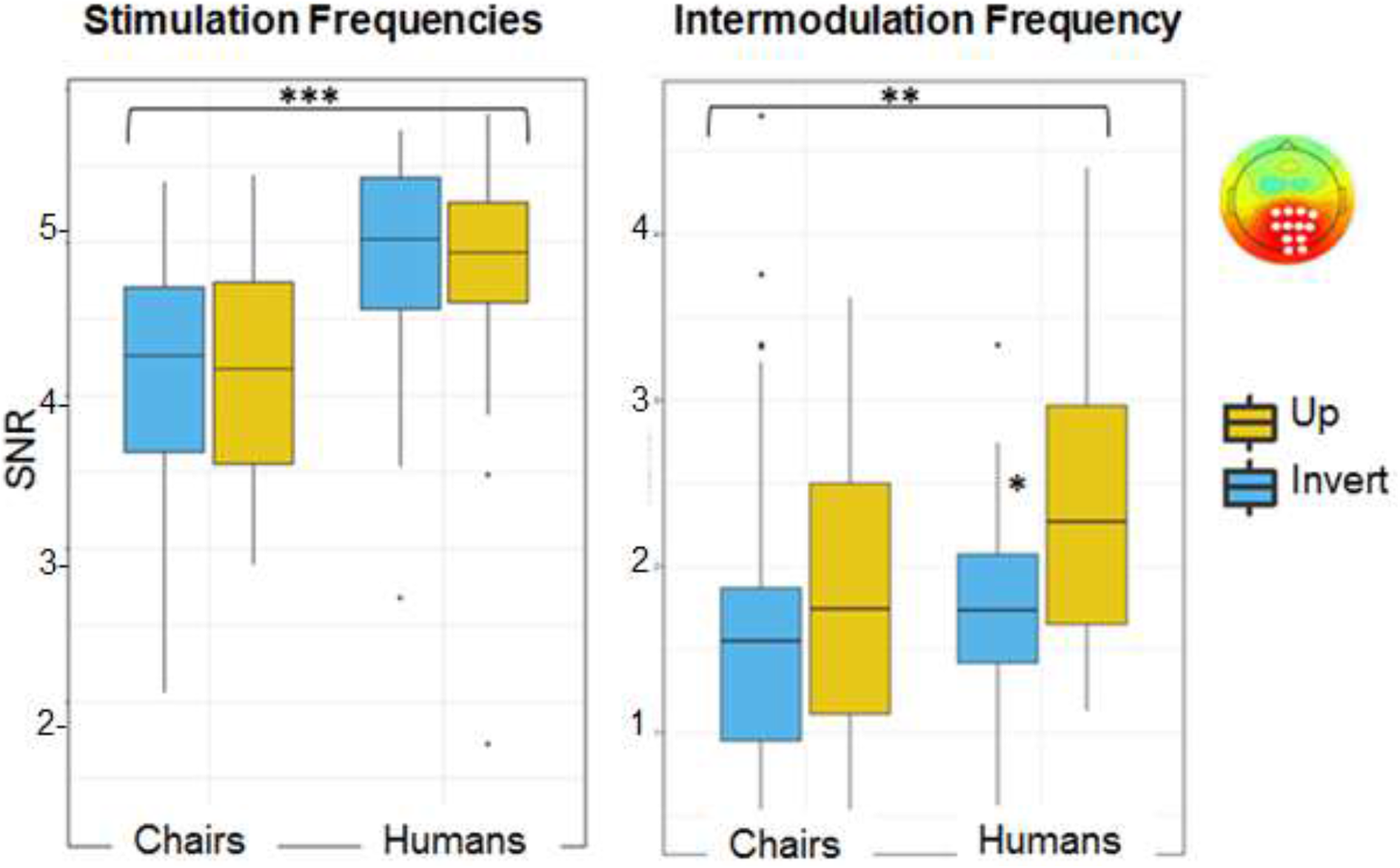
Signal to Noise Ratio averaged on participants and cortical region of analysis (white dots on topographical map) broken down by level of processing (dyad parts: stimulation frequencies, global dyad configuration: IM frequency) and type of object (Humans, Chairs). The Figure depicts inversion effect in each condition. Boxplots show median, upper and lower quartiles. Error bars represents SE. Outlying points are represented individually. Response to Humans was significantly higher than response to Chairs whatever the level of processing (dyad parts or whole). Inversion effect, however, was significant only for the whole dyad level configuration in intact upright presentation.

#### Mean first and second harmonics

Results were similar for mean f1, f2, 2f1, 2f2 SNR (Figure 4), we observed a significant main effect of Object (F (1, 29) = 13.46, p < 0. 001, η^2^ = 0.32) with a higher SNR for Humans (M = 3.52, SD = 0.57) than for Chairs: (M = 3.19, SD = 0.51), effect-size d = 0.67. There was no main effect of position (F (1, 29) < 1) nor interaction between the two factors (F (1, 29) = 1.79, p = 0.19).

In summary, there was clear response at stimulation frequencies and their harmonics to human dyad parts and so was the response to the pair of chairs parts. Only one effect was significant, the effect of Object and there was no significant interaction between Object and Position factors.

### Response at second order sum intermodulation component

For the single IM frequency component exceeding the noise threshold, f1 + f2 = 13.02, we observed a significant main effect of Object (F (1,29) = 7.07, p = 0.01, η^2^ = 0.19) with a higher SNR for Human stimuli (M = 2.12, SD = 0.88) than for Chair stimuli (M = 1.79, SD = 0.95), effect size d = 0.35. We observed also a significant main effect of position (F (1, 29) = 4.37, p = 0.04, η^2^ = 0.14) with higher SNR for upright than inverted position (UP: M = 2.14, SD = 0.98 [Humans: 2.43-1.0; Chairs: 1.86-0.89]; inverted M = 1.76, SD = 0.83 [Humans: 1.79-0.59; Chairs: 1.73-1.02]) effect size d = 0.42. There was a significant interaction between the two factors ((F (1, 29) = 4.56, p = 0.04). Figure 3 panel A shows a peak at the second order sum IM component in Human condition that is higher in Up than in Inverted position. Visual inspection of SNR topographical distribution shows that this inversion effect is the largest over right occipital cortex, as reported in previous studies using similar paradigm (e.g., Boremanse et al., 2013) and central parietal areas (Figure 5). Post-hoc comparisons confirmed that the Position effect was driven by a significantly higher response in HumanUp (M = 2.43 SD = 1.0) condition than in HumanInvert (M = 1.79, SD = 0.69) condition (t (29) = 3.16, p = 0.003, Bonferroni corrected, pcor < 0.025), effect size d = 0.75. The IM response showed similar response magnitude in ChairUp (M = 1.86, SD = 0.89) and ChairInvert (M = 1.73, SD = 1.02) conditions (t < 1).

**Figure 5.**
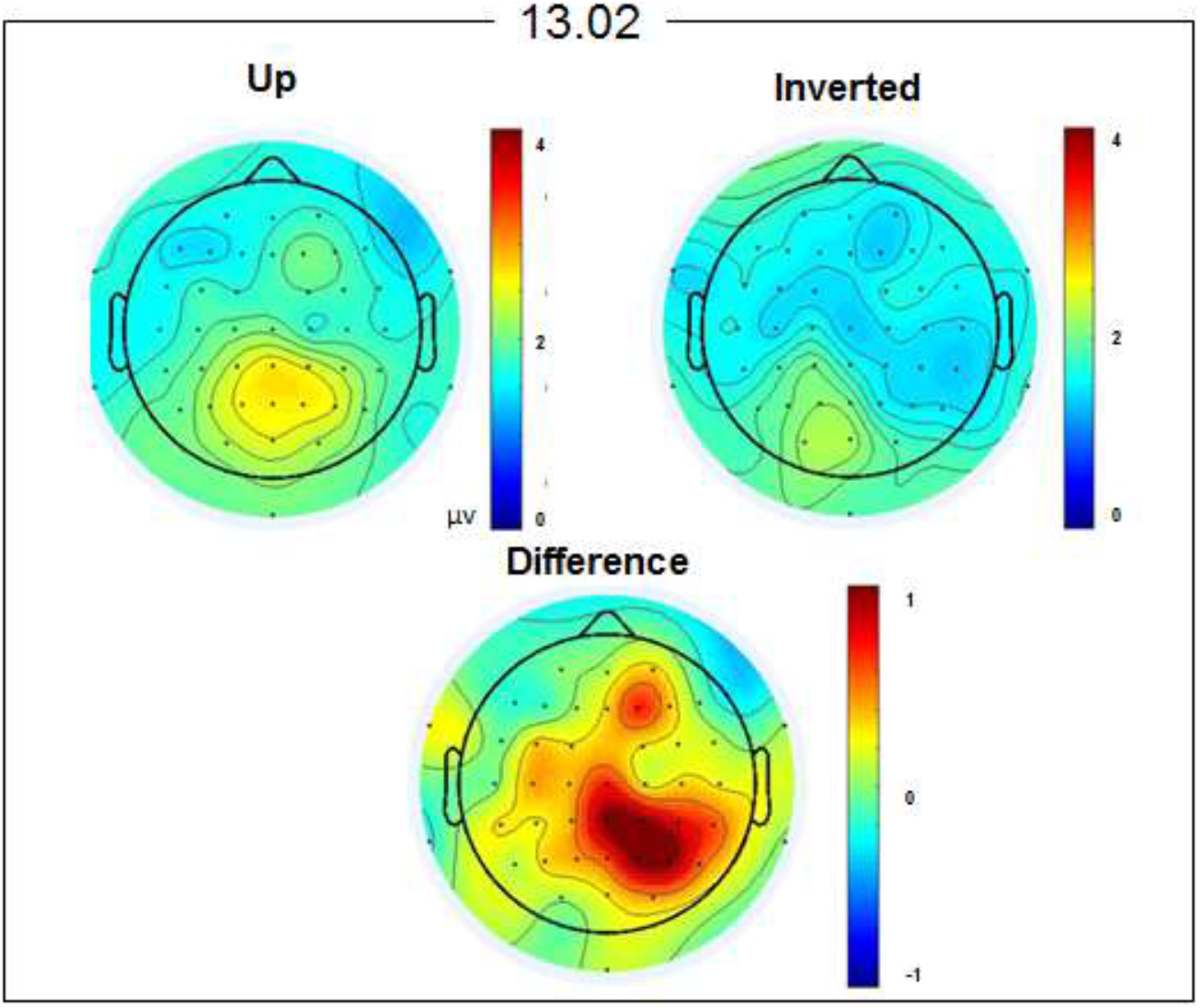
Topographical maps of SNR at intermodulation frequency f1 + f2 = 13.02 Hz (response to whole dyad configuration (average participants in Humans condition only). The figure depicts the topography of Up and Inverted conditions and their difference (inversion effect). The highest inversion effect appears over right occipito-parietal regions.

## Discussion

We hypothesized that the human visual perceptual system involves, for plurality of conspecifics, special processing that obey to gestalt principles. We studied the minimal form of a group of conspecifics, the dyad, aiming to determine whether merely perceiving two individuals in close positioning leads to perceptual grouping. Using EEG Frequency Tagging technique, we presented at two different precise frequencies each element of a dyad of silhouettes. In addition to response to parts occurring at each input frequency, we expected response to the global dyad configuration to be attested by intermodulation components. We expected inversion to impair the (holistic) response at intermodulation frequencies, but not the response to dyad parts, at stimulation frequencies. We compared the response to human dyad silhouettes to the response to silhouettes of a pair of chairs.

### Response to global dyad configuration

The critical result from the present experiment was a single IM component exceeding the noise threshold, a second order sum IM response, specific to the dyad presented in upright position. Such IM response that is not present in the input frequencies is characteristic of a particular non-linear system made of population of neurons that receives and jointly processes inputs from both dyad components (Alp et al., 2016; Appelbaum et al., 2008; Regan & Regan, 1988). Thus, our result demonstrates unambiguously that the response to the global dyad configuration is different from the response to its parts. Importantly, the IM component was significantly reduced when the dyad was presented in inverted position, an inversion effect that cannot be accounted by the response to individual silhouettes which probe to be unaffected by inverted presentation. This result supports the hypothesis of a configural processing of the dyad, driven by joint, non-linear processing of dyad parts. Hence, the present study brings neural evidence, as previously established for human faces and bodies, that the visual system processes a plurality of bodies in a configural manner, and supports the hypothesis that it is sufficient to a group of individuals to be close in space to be perceived as a coherent entity.

Moreover, IM component response to the pair of chairs was significantly reduced with respect to the response to the dyad. Note that this difference is unlikely the result of the lower number of trials in Chair condition: firstly, Chair and Human distributions are similar (Figure 4) and show comparable deviation from the mean. Secondly, overall SNR response to the pair of chairs is clearly above noise (z > 1.64, p < 0.05, on tailed) reflecting efficient processing of Chair stimuli. Even though response to chairs at intermodulation component is not null, the absence of inversion effect weakens the interpretation of a configural processing for these stimuli, and confirm previous findings that dyad inversion effect does not generalize to human-object or object-object pairs (Papeo & Abassi, 2019).

IMs can be generated by local interaction of populations of neurons whose receptive fields span separated regions of visual space tagged at different frequencies such as the border of left and right halves of a face, or figure and ground regions that abut at a boundary. In these cases, IMs has been mainly observed in medial occipital areas and almost disappeared with small physical separation (Appelbaum et al., 2008; Boremanse et al., 2013). However, when tagged stimuli are physically separated in the image, IMs, observed in more high order regions, can only arise from long-range interactions between populations of neurons that represent retinotopically distal elements (Aissani et al., 2011; Boremanse et al., 2013). In our design, stimuli were physically separated in the image. Therefore, the non-linear integration process evidenced in our data when the dyad was in upright position must be driven by a high-order structure related to a group of (two) humans in natural, common position. It will be of interest in the future to determine if similar mechanisms underlie other human plurality configurations, and to investigate the nature of these spatial high-order social structures.

Did the silhouettes position slightly leaning to each other play a role in the integration process? Our data do not allow answering this question. In Papeo et al., (2017) study, dyads of facing bodies elicited a strong inversion effect that disappeared when the exact same two figures faced away from one another. In this study, the behavioral interaction was a critical social feature. The authors concluded “Just like a pitcher with its spout leaning toward a glass, two facing bodies would be perceptually grouped by virtue of the joint action that they are engaged in”. It is important to note the difference with the present study: the dyad was not facing each other but merely present in front of the participant, either perceived as looking at him or from behind. The conclusion that we can draw from our study is that interaction between the group parts is not a necessary condition to perceive a plurality of humans as a structured unit, at least in the case of dyad. It would be interesting to investigate in more detail the spatial parameters of social configuration and to precise spatial relative positioning that is perceived as social proximity.

### Response to individual silhouettes

EEG response at stimulation frequencies f1 and f2 and their harmonics 2f1, 2f2 (response to local dyad parts), showed a highly significant effect of Object and no effect of Position, nor interaction between these factors. This result suggests that the perception of a pair of familiar objects involves the neural processing of the objects category rather than individual objects configuration. The absence of the inversion effect for human individual silhouettes seems surprising, given numerous studies having shown that body pictures and silhouettes are subjected to inversion effect (e.g., Reed et al., 2006). This pattern of perceptual processing has been previously described as the *object–inferiority effect* by (Suzuki & Cavanagh, 1995). In a visual search process they found that the global face-level representation has prior access to local-level face features. Besides, faces holistic representation, at the neural level, is also typically based on the interpretation of how the presence of an intact whole face configuration modifies the response to parts (Jacques & Rossion, 2009; Schiltz & Rossion, 2006). Our interpretation is that from two nearby bodies’ perception emerges a unitized representation that is so strong that it renders local dyad bodies constituents inaccessible to configural processing. At the cognitive level, this means that individuals are perceived differently when they are inside a dyad: individual’s body structure perception is weakened leaving room for the configural properties of the group.

## Conclusion

Recent experiments have demonstrated that dyad in face-to-face interaction is processed as a unit. These studies postulate that the relevant social feature involving specialized brain processing is the social relation between elements of the dyad, explicitly represented in the visual scene. Here, we show that perceiving two human silhouettes positioned close one from the other involves long-range communication between neural populations responding to each silhouette and triggers configural processing. This suggests that like faces and bodies, any plurality of individuals has a special perceptual status.

